# Differential regulation of lineage commitment in human and mouse primed pluripotent stem cells by NuRD

**DOI:** 10.1101/2020.02.05.935544

**Authors:** Ramy Ragheb, Sarah Gharbi, Julie Cramard, Oluwaseun Ogundele, Susan Kloet, Thomas Burgold, Michiel Vermeulen, Nicola Reynolds, Brian Hendrich

**Author notes:** Corresponding Author: Brian Hendrich, @BDH_Lab.

## Abstract

Differentiation of mammalian pluripotent cells involves large-scale changes in transcription and, among the molecules that orchestrate these changes, chromatin remodellers are essential to initiate, establish and maintain a new gene regulatory network. The NuRD complex is a highly conserved chromatin remodeller which fine-tunes gene expression in embryonic stem cells. While the function of NuRD in mouse pluripotent cells has been well defined, no study yet has defined NuRD function in human pluripotent cells. We investigated the structure and function of NuRD in human induced pluripotent stem cells (hiPSCs). Using immunoprecipitation followed by mass-spectrometry in hiPSCs and in naive or primed mouse pluripotent stem cells, we find that NuRD structure and biochemical interactors are generally conserved. Using RNA sequencing, we find that, whereas in mouse primed stem cells and in mouse naïve ES cells, NuRD is required for an appropriate level of transcriptional response to differentiation signals, hiPSCs require NuRD to initiate these responses. This difference indicates that mouse and human cells interpret and respond to induction of differentiation differently.

**Graphical Abstract:** NuRD acts like a conductor in an orchestra.
**A.** In the presence of NuRD (pink blob figure, centre) differentiation occurs in an ordered fashion in both mouse (left) and human (right) ES cells. Gene expression changes in both cell types are tightly controlled with down-regulation of pluripotency genes and up-regulation of lineage appropriate genes. This is akin to a group of musicians producing musical notes in the right order and at the right amplitude to create a coherent piece of music. **B.** Loss of “the conductor” NuRD results in increased transcriptional noise in both systems, indicated here as a low-level blanket of sound in both systems. Consequences of MBD3/NuRD loss differs between human and mouse ES cells. In mouse ES cells, differentiation cues lead to some down-regulation of pluripotency genes and incomplete progression along a lineage appropriate pathway. This is like musicians who know that they should be making music but who lose their way without a conductor’s influence. In human iPS cells the background level of noise without NuRD results in a lack of order to gene expression changes in response to differentiation. The noise from these “musicians” would be truly awful.

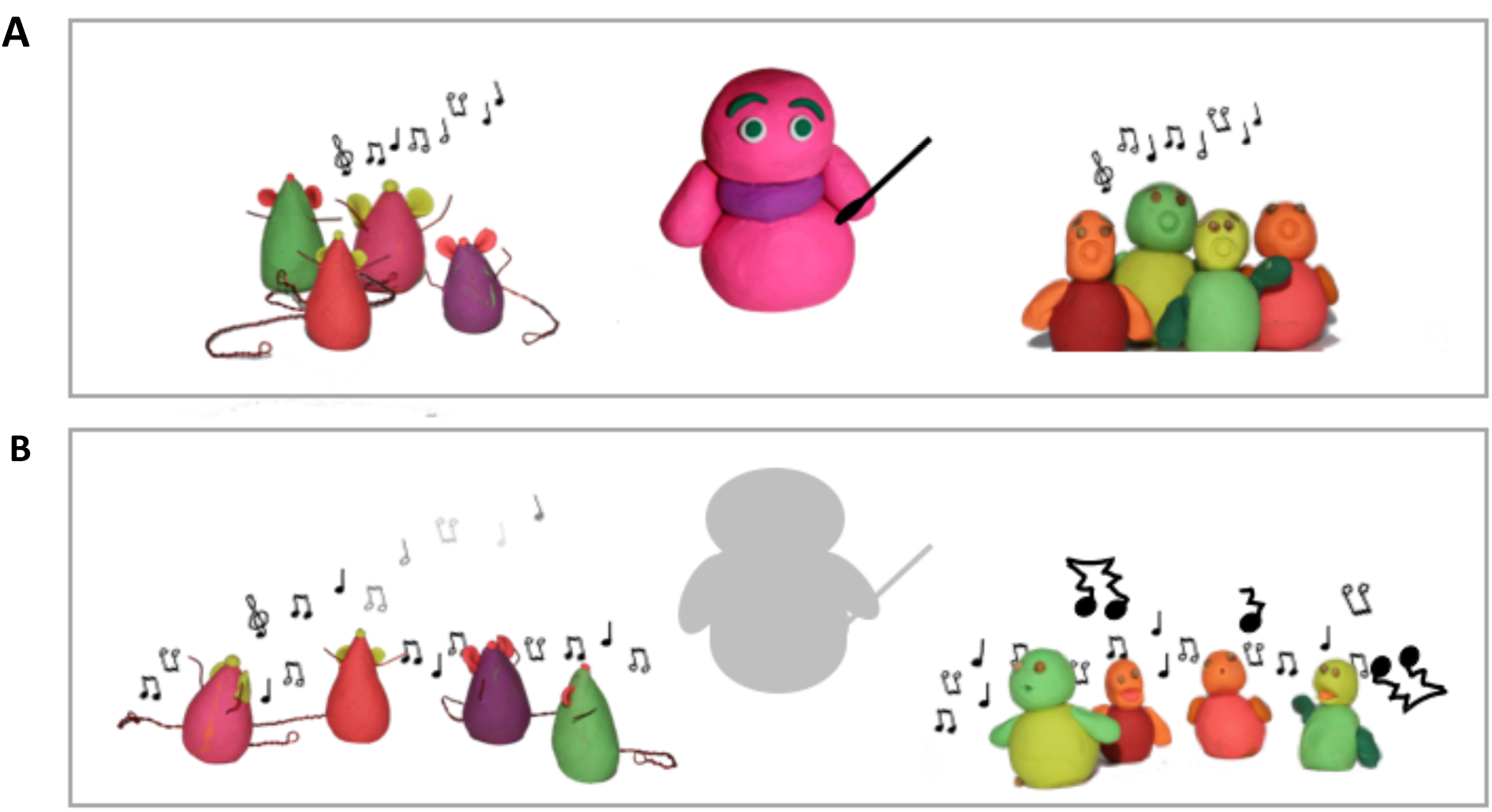

## Introduction

The identity of a eukaryotic cell is ultimately determined by its transcriptional output. The process by which cells transition from one state to another is therefore necessarily subject to tight transcriptional controls. For example, during development, in the absence of changes in external cues, transcriptional programs must remain stable for the identity of that cell to be maintained. Upon changes in external signals, transcription of some genes must be downregulated while that of others must be increased and this results in a change in cellular identity. The mechanisms which act either to maintain or change the expression state of a cell therefore underlie the ordered progression of transitions that occur throughout embryonic development. Failure of regulation of these gene expression patterns can prevent successful execution of developmental decisions, leading to developmental abnormalities, tumourigenesis or death. A comprehensive understanding of how cells control transcription during cell fate decisions is therefore critical for fields where it is desirable to control or instruct cell fate decisions, such as in regenerative medicine or cancer biology.

The ability of cells to activate or repress transcription relies largely on the conformation of the chromatin in which these genes reside. A set of chromatin remodelling complexes function to alter the structure of chromatin at regulatory elements to control gene expression (Hota & Bruneau, 2016). One such complex in particular, the NuRD (**Nu**cleosome **R**emodelling and **D**eacetylation) complex, is important for cells to undergo the changes in identity associated with the exit from pluripotency (Burgold, Barber et al., 2019, Kaji, Caballero et al., 2006, Reynolds, Latos et al., 2012a). The NuRD complex is a highly conserved multiprotein chromatin remodeller initially defined as a transcriptional repressor (Tong, Hassig et al., 1998, Wade, Jones et al., 1998, Xue, Wong et al., 1998, Zhang, LeRoy et al., 1998). NuRD activity facilitates cell fate transitions in a range of different organisms and developmental contexts (Signolet & Hendrich, 2015). The complex combines two enzymatic activities: class I lysine deacetylation, encoded by the Histone Deacetylase (Hdac) 1 and 2 proteins, with the Swi/Snf-type ATPase and nucleosome remodelling of the Chromodomain Helicase DNA binding protein 4 (Chd4). This complex also contains histone chaperone proteins Rbbp4 and 7, one of the zinc-finger proteins Gatad2a or Gatad2b, two MTA proteins (Mta1, Mta2, and/or Mta3), Cdk2ap1 and Mbd2 or Mbd3 (Allen, Wade et al., 2013, Kloet, Baymaz et al., 2015, Mohd-Sarip, Teeuwssen et al., 2017). Mbd3 is known to be required for lineage commitment of pluripotent cells and is essential for early mammalian development (Kaji et al., 2006, Kaji, Nichols et al., 2007). Structural and genetic work has found that Mbd3 physically links two biochemical and functional NuRD subcomplexes: a remodelling subcomplex containing Chd4, the Gatad2 protein and Cdk2ap1; and a histone deacetylase subcomplex containing the Hdac, the Rbbps and the Mta proteins (Burgold et al., 2019, Low, Webb et al., 2016, Zhang, Aubert et al., 2016). Mbd3 thus acts as a molecular bridge between these subcomplexes and maintains the structural integrity of NuRD.

In mouse ESCs (mESCs) Mbd3/NuRD activity modulates the transcription of pluripotency-associated genes, maintaining expression within a range that allows cells to effectively respond to differentiation signals (Bornelöv, Reynolds et al., 2018, Reynolds et al., 2012a). Despite profound developmental defects, Mbd3 deficiency in mESCs results in only moderate gene expression changes, with the majority of genes changing by less than two-fold. Rather than turning genes on or off, Mbd3/NuRD activity serves to fine-tune gene expression in mESCs (Bornelöv et al., 2018). Although this amounts to many small transcriptional changes, the cumulative effect of this is nevertheless a profound phenotype: the inability of pluripotent cells to undergo lineage commitment. While the function of Mbd3/NuRD in mouse pluripotent cells has been well defined, no study yet has defined NuRD function in human pluripotent cells.

Human and mouse ESCs can both be derived from the inner cell mass (ICM) of pre-implantation epiblasts. Yet the cell lines that emerge after culturing differ in transcriptomic, epigenetic, and morphological features (Nichols & Smith, 2009). mESCs show early developmental characteristics such as the expression of pluripotency genes, DNA hypomethylation and the activity of both X chromosomes in females. Conventional human ESCs (hESCs) are developmentally more advanced and resemble murine post-implantation epiblast or mouse epiblast stem cells (mEpiSCs), and thus are considered to be primed pluripotent (Brons et al., 2007; Tesar et al. 2007). In this study we investigated the function of the NuRD complex in human primed pluripotent stem cells (hiPSCs), and compared this to its function in mEpiSCs. We find that while MBD3/NuRD is required in both systems for cells to properly undergo lineage commitment, the way in which this function is exerted appears different. Whereas in mouse primed stem cells, as in naïve mESCs, NuRD is required for an appropriate level of transcriptional response to differentiation signals, human cells require NuRD activity to initiate these transcriptional responses. This difference in the transcriptional consequences upon loss of an orthologous protein in two different mammalian pluripotent stem cell types indicates that mouse and human cells interpret and/or respond to induction of differentiation differently.

## Results

### NuRD complex structure is conserved in mouse and human pluripotent stem cells

In order to characterise human NuRD, we used genome editing to insert coding sequence for a 3xFLAG epitope immediately upstream of the stop codon of one endogenous *MBD3* allele in human iPS cells (Fig. S1A, B). An equivalent C-terminally tagged murine endogenous MBD3 protein shows genomic localisation identical to that found for wild type MBD3 protein in mouse ES cells, and supports normal embryonic development in mice (Bornelöv et al., 2018). Biochemical isolation of MBD3/NuRD in MBD3-3xFLAG hiPSCs, or in mEpiSCs containing an identically modified *Mbd3* allele, followed by mass spectrometry identified all known components of NuRD in both systems (Fig 1A, B). A number of interacting proteins were also purified at much lower stoichiometries than was seen for core NuRD components. Comparison of mass spectrometry data between hiPSCs, mEpiSCs and mouse naïve ES cells (using MTA1-3 proteins for NuRD purification: (Burgold et al., 2019)) showed that most interacting proteins identified in human cells also interact with mouse NuRD (Fig 1C). Two cell-type specific interactors are VRTN and ZNF423, both of which are not expressed in naïve ES cells, but are found interacting with NuRD in primed PSCs (mEpiSCs and hiPSCs; Fig 1C). Two nuclear proteins were identified interacting with human NuRD that were not significantly enriched in the mouse datasets: PGBD3 and BEND3. PGBD3 is a transpose-derived protein expressed as a fusion with ERCC6 not present in mice (Newman, Bailey et al., 2008), but previously reported to interact with NuRD components in human cells (Hein, Hubner et al., 2015). Although not significantly detected in our mouse NuRD purifications, BEND3 has previously been shown to recruit NuRD to major satellite repeats in mouse cells (Saksouk, Barth et al., 2014). WDR5, ZNF296 and ZNF462 were identified interacting with mouse NuRD as described (Burgold et al., 2019, Ee, McCannell et al., 2017, Kloet, Karemaker et al., 2018), but were beneath our significance cutoff in purifications from human cells (Fig 1C). We therefore conclude that NuRD structure and biochemical interactors are generally conserved between mouse and human PSCs.

**Figure 1.**
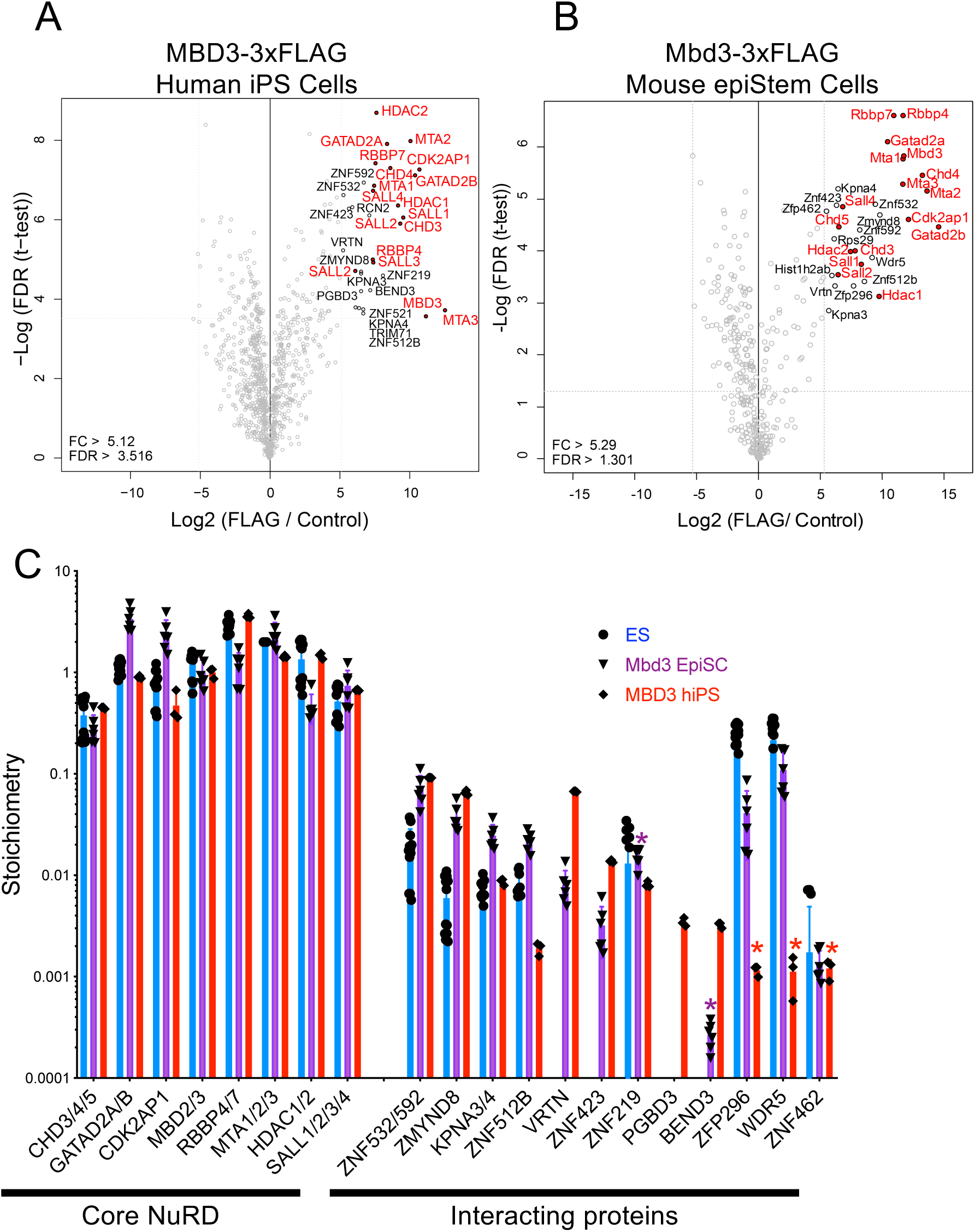
Comparison of human and mouse NuRD complexes. A) and B). Proteins associated with MBD3/NuRD in hiPSCs (A) or mEpiSCs (B) were identified by immunoprecipitation and mass spectrometry. Proteins significantly associating with MBD3 are indicated, with NuRD component proteins indicated in red. The human data comprise three independent immunoprecipitations from one preparation of nuclear extract, while the mouse data comprise three independent immunoprecipitations each from nuclear extract preparations made from two independent cell lines. C) Relative enrichment of indicated proteins normalised to the bait protein for each experiment: for mouse, ES cell data (taken from (Burgold et al., 2019); blue) were normalised to two MTA proteins, while for both the epiSC (purple) and hiPS (red) experiments the data were normalised to one MBD3 protein. Error bars represent standard deviation of three (hiPSCs), six (mEpiSCs) or nine (mESCs) replicates. Asterisks indicate situations where the protein enrichment was not significant in this cell type, but the stoichiometry is displayed for comparison.

We next asked what function was played by MBD3/NuRD in human PSCs. To this end we used CRISPR/Cas9-mediated gene targeting to create an *MBD3*-KO iPSC line (Fig. 2A; Fig S1C). Immunoprecipitation with the remodelling subunit of NuRD, CHD4, allowed for purification of other complex components in wild type cells, but not in the *MBD3*-KO cells, indicating that human NuRD does not form without MBD3 (Fig 2B). Interactions between CHD4 and other NuRD components were restored when an MBD3 transgene was overexpressed in the null cells (“Rescue”, Fig 2A,B), indicating that the transgenic MBD3 is sufficient for NuRD formation as it is in mESCs (Bornelöv et al., 2018, Reynolds, Salmon-Divon et al., 2012b).

**Figure 2.**
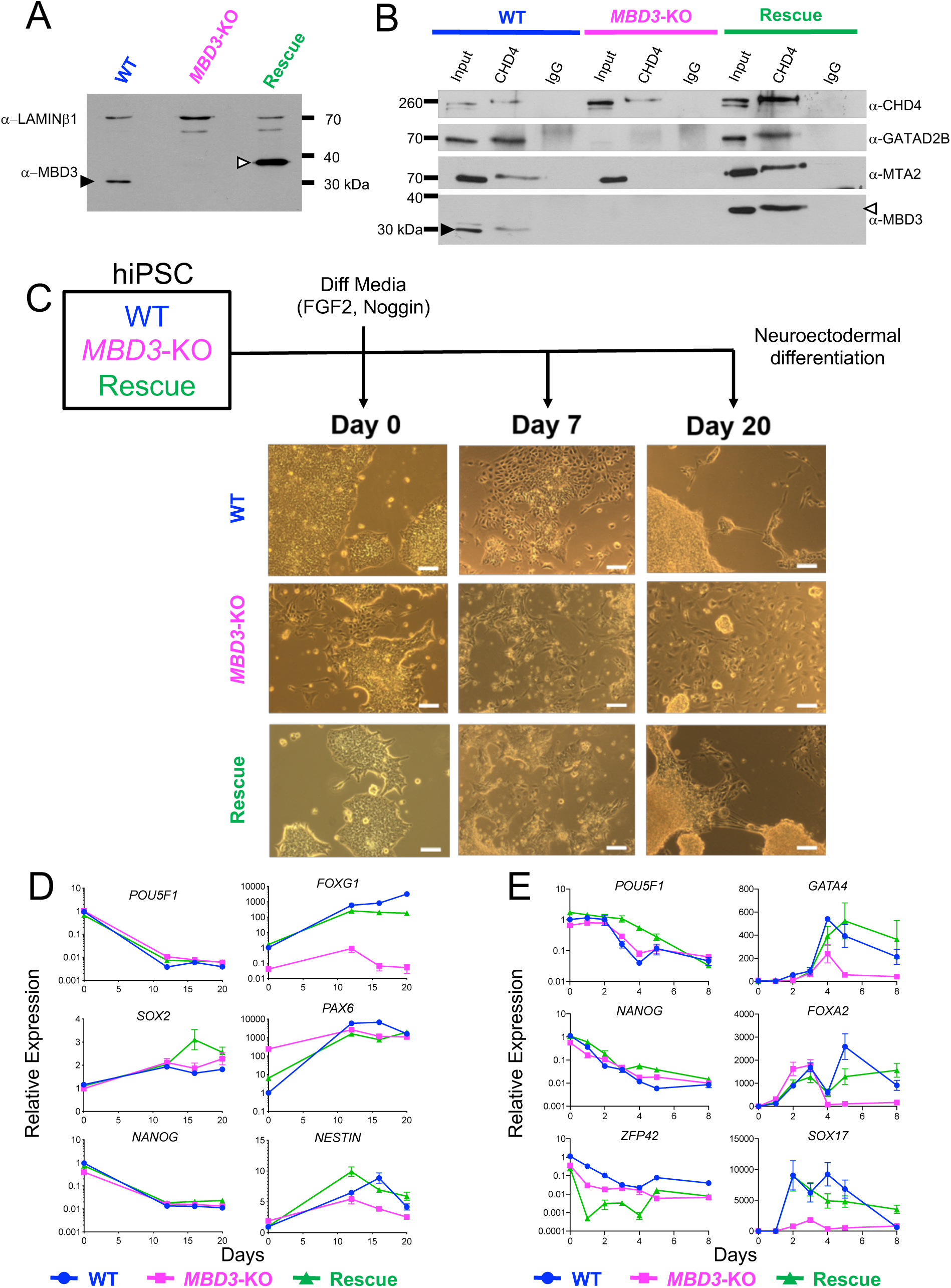
Human iPS cells lacking MBD3/NuRD fail to undergo programmed differentiation. A) Western blot of nuclear extracts from wild type (WT), *MBD3*-KO and *MBD3*-KO hiPSCs rescued with an MBD3-3xFLAG transgene (“Rescue”). The blot was probed with antibodies indicated at left. The closed arrowhead indicates native MBD3, while the open arrowhead indicates the MBD3-3xFLAG fusion protein. B) Nuclear extracts from wild type, *MBD3*-KO and Rescued cells was immunoprecipitated with anti-Chd4, western blotted and probed with antibodies indicated at right. Arrowheads as in Panel A. C) Scheme of the differentiation experiment (top) and images of indicated cell cultures at Day 0, 7 or 20 of differentiation. Scale bar indicates 100 µm. D) Expression of pluripotency (*POU5F1, SOX2* and *NANOG*) and lineage specific genes (*FOXG1, PAX6* and *NESTIN*) during neural differentiation was measured by qRT-PCR. Y-axis represents expression relative to that in wild type cells at Day 0, while the X-axis represents the time in days. Error bars represent the standard deviation of ≥3 biological replicates. E) as in panel D, but for definitive endoderm differentiation protocol. Pluripotency-associated genes (*POU5F1, NANOG, ZFP42*) on the left, and differentiation-associated genes (*GATA4, FOXA2, SOX17*) at right.

*MBD3*-KO hiPSCs were viable in standard culture conditions (mTESR or E8 (Chen, Gulbranson et al., 2011)), though unlike wild type cells null cultures showed some degree of spontaneous differentiation in both culture conditions (Fig 2C). While wild type and Rescue cultures presented as morphologically homogeneous colonies with clear boundaries, mutant cultures were mix of cells showing a compact, undifferentiated morphology as well as a population of flatter, less dense colonies with irregular boundaries, reminiscent of differentiated cells (Fig. 2C). This was surprising since *Mbd3*-KO mESCs are resistant to differentiation (Kaji et al., 2006).

To determine whether human NuRD is required for lineage commitment of pluripotent cells, wild type, mutant and Rescued hiPSCs were induced to differentiate towards a neuroectodermal fate or a definitive endoderm fate (see Methods). After 20 days of neuroectodermal differentiation, axon-like extensions were readily identifiable in wild type and Rescue cultures (Fig. 2C). In contrast, no such appendages were found in *MBD3*-KO cultures, indicating a requirement for MBD3/NuRD for successful completion of this differentiation process. All three cell lines showed a decrease in expression of pluripotency markers across both differentiation protocols, indicating that NuRD is not required for PSC to respond to differentiation signals (Fig. 2D, E). *MBD3*-KO cells failed to properly induce expression of some lineage-appropriate genes in both differentiation protocols, but this ability was restored upon rescue with the MBD3 transgene (Fig. 2D, E). While NuRD is therefore required in human cells to faithfully maintain a self-renewing state, it is also required for appropriate lineage determination in these two differentiation protocols.

### NuRD mutant hiPS cells are unable to maintain a stable pluripotency state

To determine how NuRD facilitates lineage commitment in hiPSCs, we analysed and compared the transcriptomes of WT, *MBD3*-KO and Rescue cells at 0, 3, 6 and 12 days upon neuroectodermal differentiation. Visualising the data using a multidimensional scaling plot (Ritchie, et al., 2015) separated each sample along the time of differentiation, represented by PC1, and the genotype, represented by PC2 (Fig. 3A). Data from WT and Rescue cells clustered close to each other and followed a similar developmental trajectory, indicating that overexpression of *MBD3* does not dramatically impair early stages of neural differentiation. In contrast, data from *MBD3*-KO cells clustered separately, indicating that they are undergoing aberrant differentiation, consistent with our RT-qPCR data (Fig. 2D). At day 0 NuRD mutant cells occupy a position further along the differentiation trajectory (PC1) than do either WT or Rescue cells, likely resulting from the presence of morphologically differentiated cells within the self-renewing *MBD3*-KO cultures (Fig. 2C).

**Figure 3.**
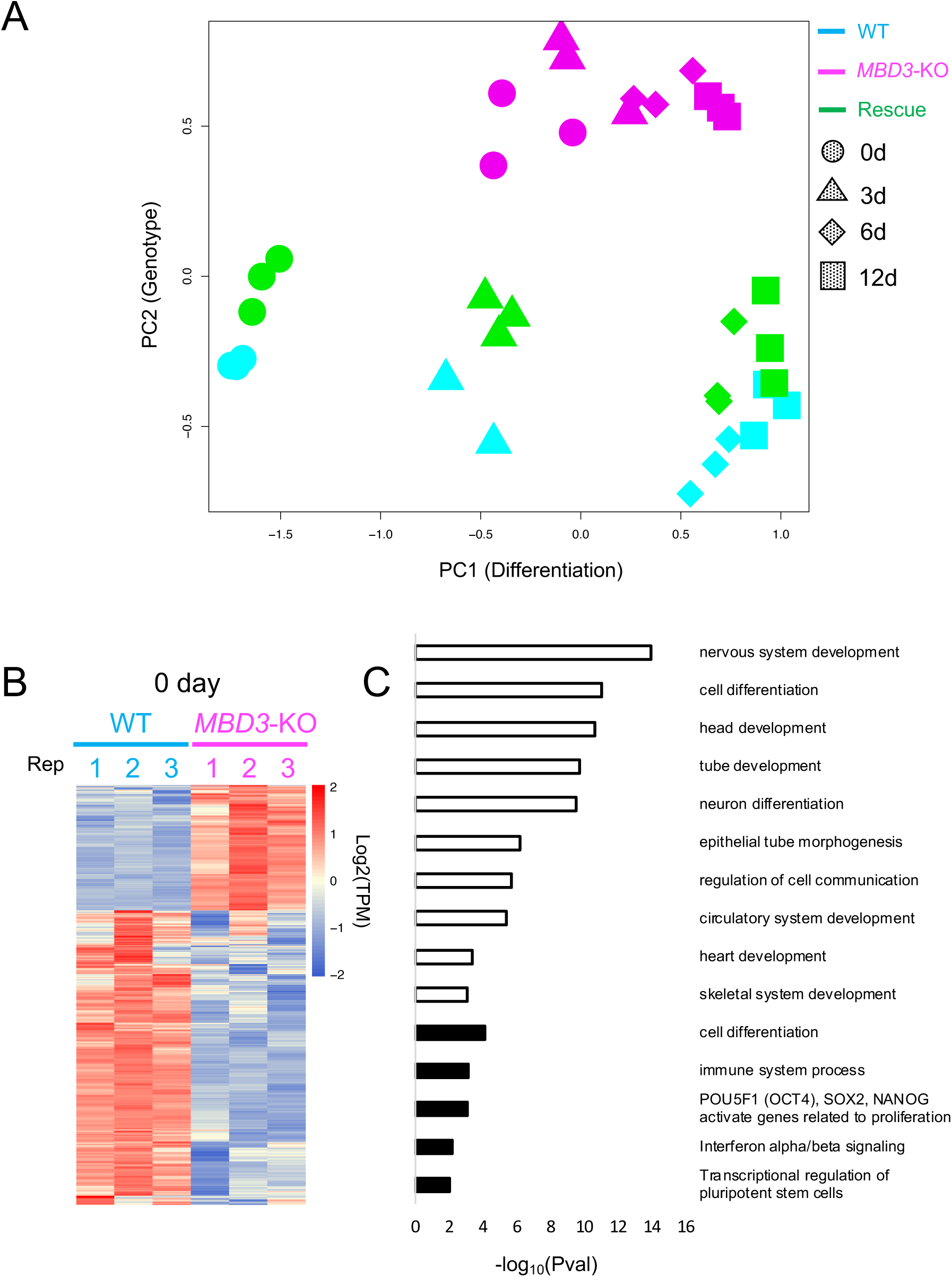
NuRD is required to maintain a stable pluripotency state in hiPSCs. A) MDS plot made from RNA-seq data of wild type, *MBD3*-KO and Rescued hiPSCs across a neural differentiation timecourse. Each point represents a biological replicate, and shapes indicate the days of differentiation. B) Heat map of genes found to be differentially expressed between WT and *MBD3*-KO cells in self-renewing conditions (day 0; FDR 5%). C) GO terms associated with genes significantly activated (open bars) or repressed (filled bars) in *MBD3*-KO cells relative to WT hiPSCs in self-renewing conditions.

To try to understand why *MBD3*-KO PSC were unable to stably maintain an undifferentiated state, the transcriptomes of WT and *MBD3*-KO PSCs were compared in self-renewing conditions (Fig. 3B). Null cells showed 823 differentially expressed genes compared to wild type cells (246 up- and 577 down-regulated, FDR 1%). GO term enrichments performed on the set of down-regulated genes showed terms related to pluripotency (Fig 3C, Table S1) consistent with a failure to maintain a stable pluripotent state in the absence of MBD3/NuRD. Expression profiles of pluripotency markers (*NANOG, TDGF1, FOXD3* and *FGF2*) showed a significant down-regulation in mutant cultures when compared to the WT or Rescue cells (Table S1). The 246 up-regulated genes showed enrichment for terms related to the development of different lineages (Fig 3C, Table S1). Given that *MBD3*-KO cells failed to undergo programmed neural or endodermal differentiation despite precociously expressing differentiation markers, we conclude that, like in mESCs (Burgold et al., 2019), human NuRD functions to prevent inappropriate gene expression in undifferentiated pluripotent cells, and this noise reduction function is important for faithful execution of lineage decisions.

### Human NuRD activity is required for appropriate transcriptional response to differentiation signals

To better understand how human NuRD facilitates lineage commitment, we next asked how gene expression changes during the differentiation time course differed in *MBD3*-KO cells as compared to WT or Rescue cells. By considering both genotype and the time of differentiation in our differential analysis, we identified genes showing expression changes in at least one cell line compared to the others during differentiation. Clusters of co-expressed genes were identified using K-means clustering (Soukas, Cohen, Socci, & Friedman, 2000), resulting in 6 groups showing similar expression profiles (Figure 4A).

**Figure 4.**
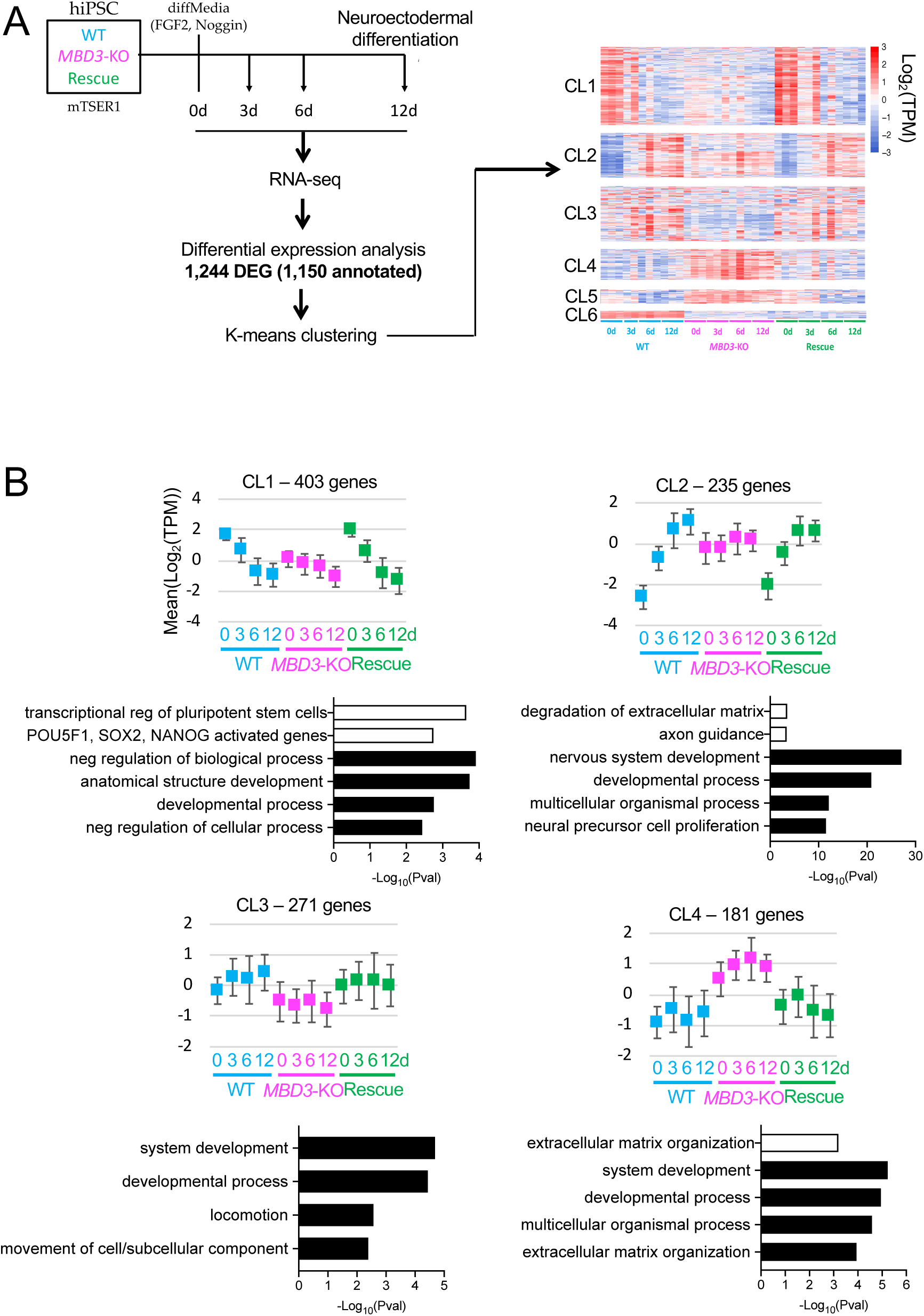
NuRD is required for transcriptional responsiveness in hiPSCs. A) Scheme of the experiment. Wild type (WT), *MBD3*-KO or Rescued cells maintained in mTESR1 media were subjected to neural differentiation. Cultures were sampled at indicated time points for RNA-seq, leading to the identification of 1150 annotated differentially expressed genes (DEG; FDR 1%). Kmeans clustering of genes by expression pattern led to the heat map shown at right, with six major gene clusters. B) Mean expression for genes in each cluster is displayed across the differentiation time course for each cell line. Error bars indicate standard deviations of average expression. The four most significant GO terms associated with each cluster are plotted as solid bars, while up to two pathways (P.adj≤0.01) are also plotted in open bars. A full list of GO terms and pathways is available in Table S4 and S5.

Cluster 1 is composed of 403 genes down-regulated during normal differentiation (Fig. 4A, B; Table S2, S3 and S4). These genes were generally underexpressed in *MBD3*-KO hiPSCs, yet become further down-regulated as cells are subjected to differentiation conditions. This cluster includes pluripotency-associated genes such as *POU5F1, NANOG, FOXD3, TDGF1, FGF2, ZSCAN10, DPPA4* and *PRDM14*, validating and extending our conclusion drawn from data shown in Fig. 3 that *MBD3*-KO hiPSCs have a defect in maintaining the self-renewing state in standard conditions. Transcription factor binding sites enriched within 10Kb of the TSS of genes in this cluster showed significant enrichment of consensus binding sites for general transcription factors associated with activation of pluripotency gene expression (i.e. MYC, ATF, CEBPB; Table 1), consistent with a decrease in expression of pluripotency-associated genes. This analysis additionally identified consensus binding sites for SNAIL and ZEB1 (Table 1), transcription factors associated with epithelial to mesenchymal transition as well as repression of pluripotency gene expression (Jiang, Yan et al., 2018, Moreno-Bueno, Portillo et al., 2008). Human ES cells undergo EMT as part of the differentiation process (Kim, Yi et al., 2014), so this likely results from inappropriate expression of epithelial genes which would normally precede an EMT event.

**Table 1:**
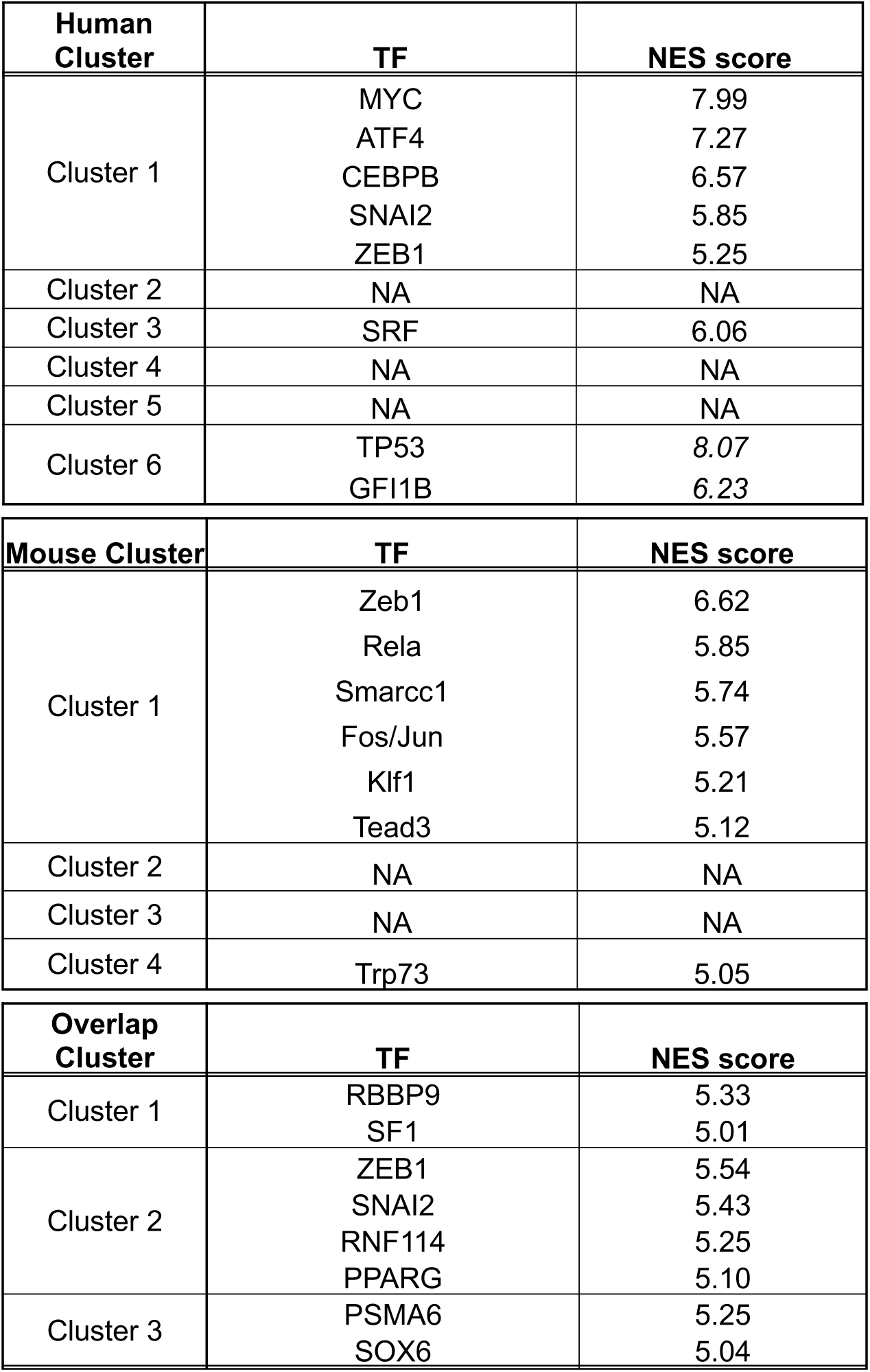
Transcription factor binding sites associated with gene clusters.

Genes in clusters 2-4 are predominantly associated with GO terms involved in differentiation (Fig. 4B; Table S2, S3 and S4). Cluster 2 contains genes associated with neuroectodermal differentiation and were induced in wild type and Rescue cells. While Cluster 2 genes (including *PAX6, OTX1* and *SOX1)* showed inappropriate expression in *MBD3*-KO cells at time 0 and remained high throughout the differentiation time course, Cluster 3 genes, some of which are associated with neuronal maturation (such as *PLP1, SEMA3A*, and *APP*) were expressed at a lower level in mutant cells than in either WT or Rescue cells. Cluster 4 contains genes not induced in either WT or Rescue cells, but which showed inappropriate expression in *MBD3*-KO cells at all time points (e.g. *WNT5A, FOXA2, PAX7*; Fig 4B, Table S2). Cluster 5 contains only 27 genes which fail to be appropriately silenced during differentiation in *MBD3*-KO cells, but show no significant enrichment with any GO term, and genes in Cluster 6 show similar expression patterns in mutant and Rescue cells, and are hence unlikely to contribute to the differentiation failure phenotype of *MBD3*-KO cells (Fig. S2). Genes in Clusters 2 and 4 are not associated with any specific TF binding sites, while Cluster 3 genes show enrichment of binding sites for the general transcription factor SRF (Table 1). This lack of evidence for misregulation of a specific transcriptional programme indicates that *MBD3*-KO hiPSCs fail to interpret a range of different differentiation signals, as opposed to just one or two main pathways. The silencing of pluripotency-associated genes (Cluster 1) in *MBD3*-KO cells demonstrates an ability of these cells to respond to the loss of self-renewal signals, yet the failure of genes in Clusters 2 and 5 to appropriately change expression during the differentiation time course indicates that NuRD is required for cells to properly respond to differentiation signals.

### MBD3/NuRD controls lineage commitment differently in human and mouse primed PSC

Mbd3/NuRD facilitates exit from the self-renewing state in mouse naïve ESCs (Burgold et al., 2019, Kaji et al., 2006) and thus it was surprising that human primed PSC required MBD3/NuRD to properly maintain the self-renewing state. To determine whether this was a difference between human and mouse cells, or between naïve and primed pluripotent cells we next asked whether NuRD was required to maintain the self-renewing state in mEpiSCs. *Mbd3*^*-/-*^ mEpiSCs were derived in culture from *Mbd3*^*-/-*^ mESCs or by transient Cre expression in mEpiSCs derived from *Mbd3*^*Flox/-*^ mESCs (See Methods). *Mbd3*^*-/-*^ mEpiSCs derived through either method were indistinguishable, and appeared uniformly undifferentiated (Fig. S3A), indicating that the spontaneous differentiation seen in hiPSCs does not reflect a general requirement for MBD3/NuRD to maintain primed PSC.

To further compare mouse and human primed PSC, gene expression was monitored across a neural differentiation time course in both Floxed and *Mbd3*^*-/-*^ mEpiSCs by RNA-seq (Fig. 5A; Table S5, S6 and S7). When the data are visualised using a multidimensional scaling plot (Chen, Lun, & Smyth, 2016), each sample separates along PC1, representing time of differentiation, and PC2, representing genotype (Fig 5B). Control and *Mbd3*^*-/-*^ cells occupy the same position along the differentiation trajectory (PC1), consistent with our observation that mEpiSCs do not require MBD3/NuRD to maintain a morphologically undifferentiated state.

**Figure 5.**
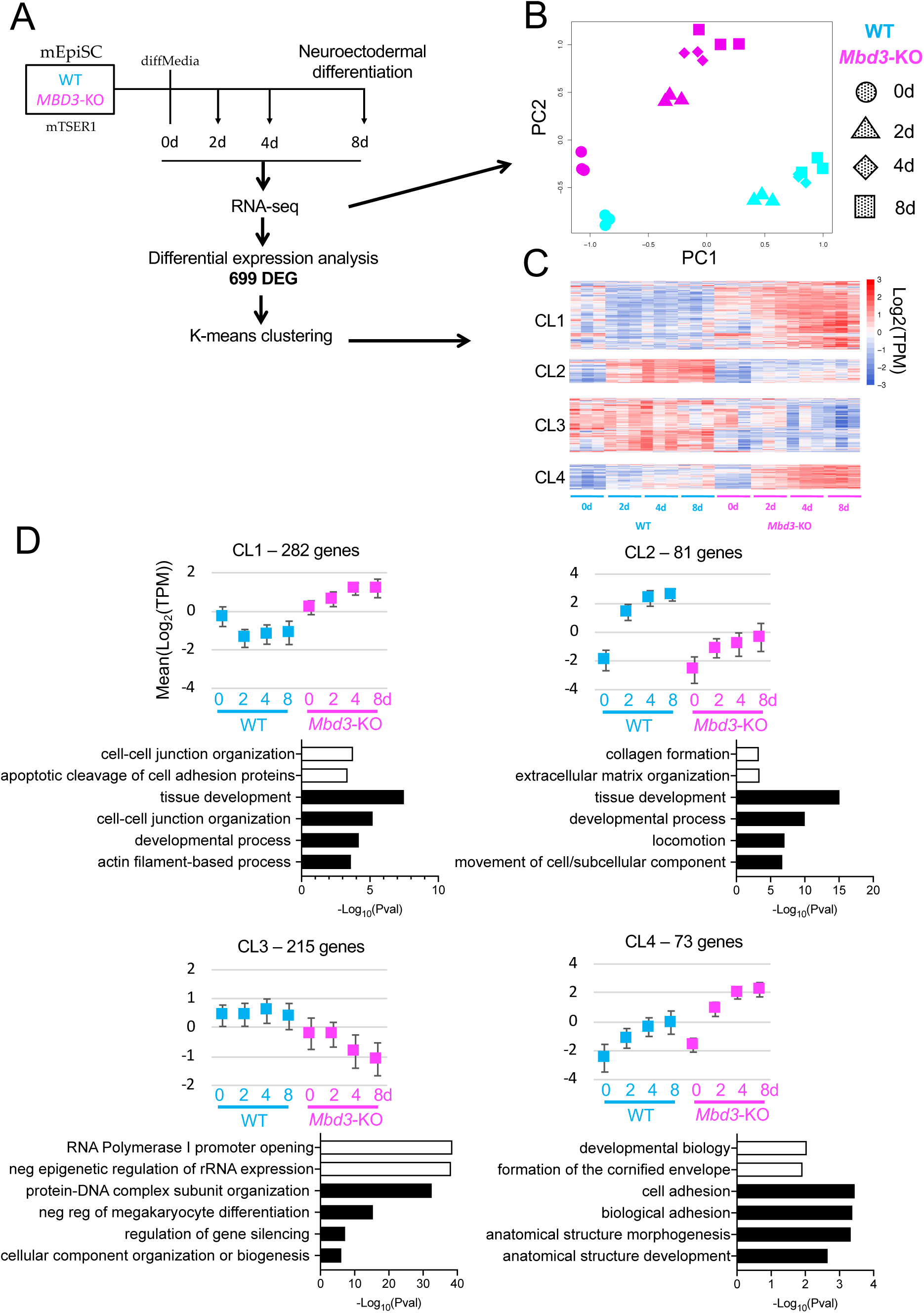
NuRD facilitates an appropriate transcriptional response in mEpiSCs. A) An outline of the experiment, as in Figure 4. B) MDS plot of gene expression data collected across the neural differentiation time course, as in Figure 3A. C) Heat map of DEG (FDR 1%) separated into four clusters by K-means clustering. D) Mean expression and most significant GO terms for each cluster as in Figure 4B. A full list of GO terms and pathways is available in Table S6 and S7, and expression plots and GO terms associated with clusters 5 and 6 are shown in Fig. S2.

As with the hiPSCs, we considered both the genotype and the time of differentiation in our analysis and thus identified 699 differentially expressed genes. Co-expressed genes were grouped using K-means clustering, resulting in 4 clusters showing similar expression profiles (Fig. 5C). In contrast to the human cells which generally showed a lack of transcriptional response to the differentiation time course, clusters identified in mouse cells showed transcriptional responses, but these responses differed from those in wild type cells (Fig. 5D). Cluster 1 genes showed decreased expression in wild type cells across the time course, but increased expression in *Mbd3*^*-/-*^ cells. Genes in clusters 2 and 4 also show increases in both wild type and mutant cells, but Cluster 2 genes showed a reduced response in mutant cells, whereas cluster 4 genes showed an increased response. Cluster 3 genes decreased in expression in mutant cells across the time course, but showed no overall change in wild type cells. In all clusters the genes are associated with very general GO terms, and are unlikely to represent individual pathways or developmental trajectories (Fig. 5D; Table S6 and S7). These data indicate that in mouse EpiSC Mbd3/NuRD is not strictly required for cells to respond to differentiation signals as was seen in hPSC, but rather is required for an appropriate level of response, as it is in naïve mESC (Burgold et al., 2019). Rather than being required for the transcriptional response to differentiation cues as in the human differentiation course, Mbd3/NuRD functions in mouse primed PSC to facilitate an appropriate transcriptional response to neural induction.

Despite the differences in NuRD-dependent gene regulation observed in mouse and human primed PSC described thus far, we asked whether there could be a conserved core set of genes regulated similarly in primed PSC from both species, which might contribute to the shared requirement for NuRD in lineage commitment. Comparing gene expression datasets between the human and mouse experiments (Fig. 6A) identified 153 genes, the orthologues of which were differentially expressed in both human and mouse cells. K-means clustering using expression data from the human cells segregated the genes into three main clusters (Fig. 6; Table S8). Orthologues defined in human clusters showed 3-4 different gene expression profiles in mouse cells (Table S10), generally showing a reduced or inappropriate transcriptional response in the absence of Mbd3/NuRD (Fig. 6). One exception is a subset of thirteen Cluster 2 genes (excluding *Mbd3*) which are rapidly activated in wild type cells but fail to be activated in mutant cells across the time course (human Cluster 2; corresponding mouse subcluster 1: Fig. 6; Table S10). Notably these genes are underexpressed in mutant mEpiSCs in self-renewing conditions, but overexpressed in hiPSCs, indicating that they are not regulated similarly in the two cell types. We therefore conclude that the transcriptional consequences of Mbd3/NuRD loss are different in human versus mouse primed PSC, but in both cases this activity is required for cells to properly undergo lineage commitment.

**Figure 6.**
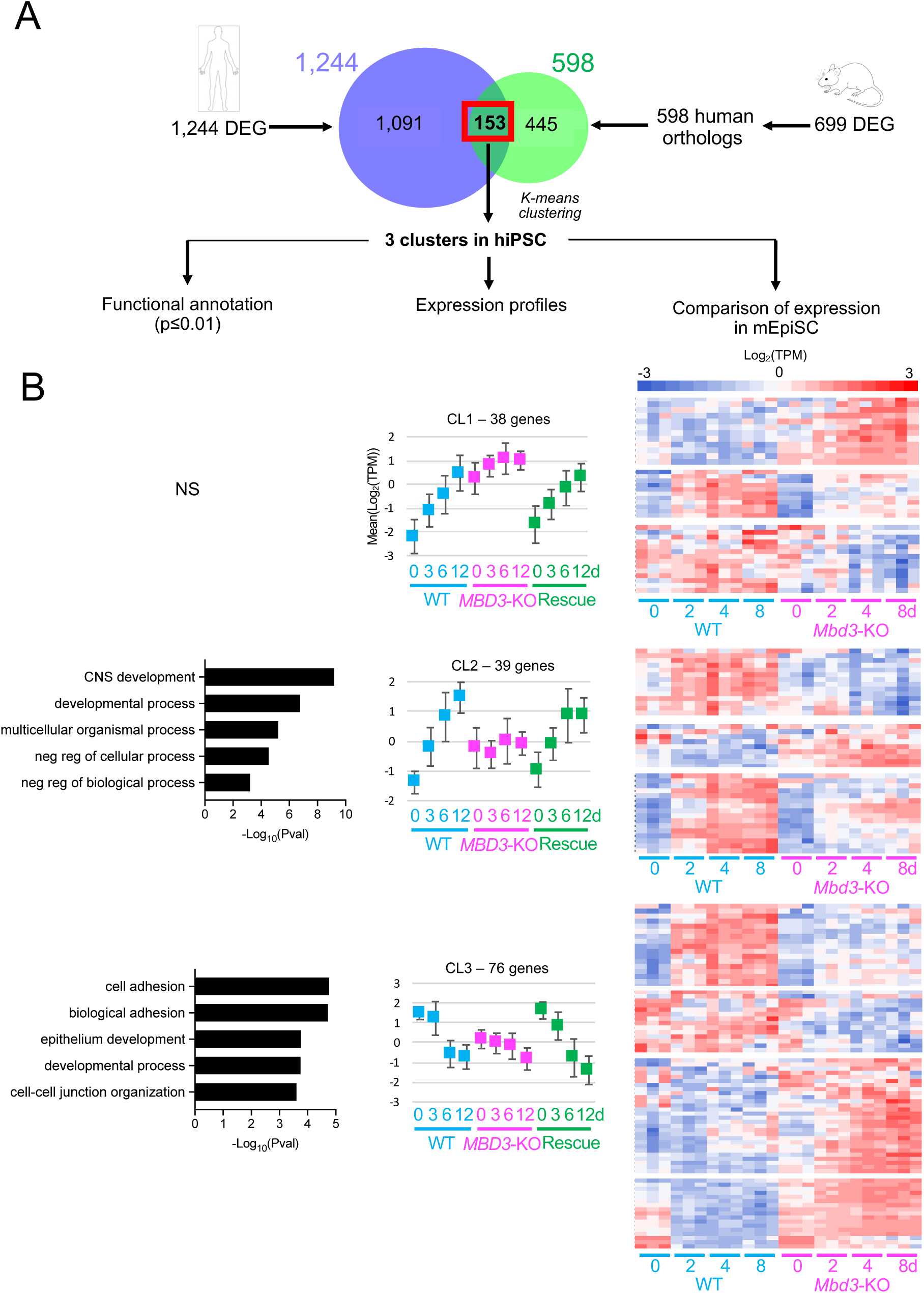
MBD3/NuRD deficiency elicits a different response in human and mouse primed pluripotent stem cells. A) Schematic of the experiment: the Venn diagram shows the overlap of differentially expressed genes identified in human cells, and the identified human orthologues of those identified in mEpiSCs. K-means clustering of this set of genes in human data led to the formation of three gene clusters. B). GO terms (“Functional annotation”; P.adj ≤ 0.01) and expression profiles are shown for the human gene clusters. Error bars indicate standard deviations of average expression. K-means clustering has been performed on the mouse gene orthologues of each human cluster resulting in 3 subclusters related to human cluster 1, 3 subclusters related to human cluster 2 and 4 subclusters related to human cluster 3. Heat maps of expression for orthologues in the mouse differentiation experiment are shown at right. There were no significant GO terms associated with Cluster 1 genes.

## Discussion

Differentiation of mammalian pluripotent cells involves large-scale changes in transcription, which result in loss of one cell identity and gain of a new, more differentiated identity. Orchestrating these changes in transcription are a large cast of different transcription factors and signalling molecules, but there is also a set of chromatin remodellers whose activity is essential to initiate, establish and maintain a new gene regulatory network (GRN) (Gokbuget & Blelloch, 2019, Hota & Bruneau, 2016). When induced to differentiate, both mouse and human pluripotent stem cells depend on NuRD activity to elicit an appropriate transcriptional response and undergo lineage commitment, but the manner in which NuRD is used to facilitate this response differs. Induction to differentiate elicits changes in transcription from a range of genes in both human and mouse PSC. The absence of MBD3/NuRD activity in human cells results in a subset of these genes failing to respond to the differentiation cues, while in mutant mouse PSC the response is present and widespread, but often muted or inappropriate (Figs 4, 5). NuRD activity is additionally required to maintain the pluripotency GRN of hiPSCs cultured in self-renewing conditions, but neither primed nor naïve mouse PSCs display this requirement (Fig. S3A and (Kaji et al., 2006)). We see no large-scale differences in the biochemical make-up of NuRD between human and mouse primed stem cells, which would be consistent with the human and mouse complexes exerting similar, or identical biochemical functions. The observed differences in the consequences of MBD3 deficiency are therefore likely to result from subtle differences in how NuRD activity is used by the cells to respond to changes in environment.

One example of how human and mouse cells respond differently to loss of MBD3/NuRD is in regulation of the *ZEB1/Zeb1* genes. *ZEB1* is overexpressed in self-renewing human PSC and remains high through the neural induction time course, whereas in mouse cells *Zeb1* is underexpressed in self-renewing cells and fails to be activated during the time course (Fig. S4). ZEB1 has been shown to repress polarity and gap-junction genes associated with an epithelial morphology, promoting an epithelial to mesenchymal transition (EMT) (Aigner, Dampier et al., 2007). This function is required for neural differentiation in vivo and from hESCs in culture (Jiang et al., 2018, Singh, Howell et al., 2016). It is not surprising, then, that Human Cluster 1 genes, which show underexpression throughout the differentiation time course in mutant cells and show enrichment for cell adhesion genes (p = 2×10^−3^; Table S4 and S5), are also enriched for ZEB1 DNA binding motifs (Fig. 4B; Table 1). In mouse cells, however, Zeb1 motifs were associated with the cluster of genes highly associated with cell-cell junctions and showing inappropriately high expression levels at all time points (Cluster 1: Fig. 5D and Table 1), consistent with aberrantly low expression of the Zeb1 repressor. While transgenic overexpression of ZEB1 was reported to increase neural differentiation of hESCs (Jiang et al., 2018), it did not lead to precocious differentiation of self-renewing hESCs, and hence is unlikely to be the principal factor behind the precocious differentiation seen in hiPSCs lacking MBD3.

It is possible that differences in transcriptional responses to differentiation in human and mouse cells could be due to the fact that, unlike mouse cells, human PSC are unable to maintain a stable self-renewing state in the absence of MBD3/NuRD, and are, in effect, responding to loss of self-renewal conditions when they have already started to differentiate. One possible, trivial explanation for this difference in the ability of mouse and human PSC to self-renew could be due to differences in the constituents of media used for self-renewal culture. Both mEpiSC culture and hiPSC culture rely on FGF2 and activation of SMAD2/3 through addition of Activin or TGFβ (Brons, Smithers et al., 2007, Chen et al., 2011, Tesar, Chenoweth et al., 2007), while naïve mouse ES cells are maintained through LIF signalling and dual inhibition of GSK3 and MEK/ERK (Ying, Wray et al., 2008). One consistent difference between mouse and human PSC culture media is the inclusion in human media of ascorbic acid (Vitamin C). Ascorbic acid has been shown to increase the activity of TET enzymes, which promote the demethylation of 5-methylCytosine in DNA, though this has been shown to promote a more naïve state, rather than promote differentiation (Blaschke, Ebata et al., 2013, Yin, Mao et al., 2013). *Mbd3*-KO mESCs contain a reduced amount of DNA methylation relative to wild type cells (Latos, Helliwell et al., 2012), consistent with them being less able to differentiate. It is therefore unlikely that an increase in TET enzyme activity would be behind the precocious differentiation seen in hPSC cultures. Rather, we suggest that the differences observed between human and mouse PSC in self-renewal or the ability to initiate an appropriate developmental response are most likely due to differences between the two species. As pointed out previously (Takashima, Guo et al., 2014), primates have not evolved the ability to undergo embryonic diapause (Nichols & Smith, 2012), and hence pluripotency may be a less stable state in humans than in mice, and consequently be less tolerant to the loss of a major chromatin remodelling complex such as MBD3/NuRD.

The mouse and human NuRD complexes present in primed PSC appear to be biochemically very similar, and our methods identified no notable species-specific interactors or alternate stoichiometries (Fig. 1). NuRD is found at all active enhancers and promoters in both mouse and human cells (Bornelöv et al., 2018, Burgold et al., 2019, de Dieuleveult, Yen et al., 2016, Gunther, Rust et al., 2013, Miller, Ralser et al., 2016, Shimbo, Du et al., 2013), but only a relatively small proportion of these genes changes expression after *MBD3* deletion (Figs. 4A, 5A) (Bornelöv et al., 2018). This is because NuRD acts to fine-tune expression through nucleosome mobilisation, and to cement longer-term gene expression changes through histone deacetylation activity (Bornelöv et al., 2018, Liang, Brown et al., 2017). NuRD’s fine-tuning function also works to ensure cells are able to respond appropriately when stem cells are induced to undergo lineage commitment. Yet the actual series of molecular events through which chromatin remodellers facilitate a cell’s ability to respond to differentiation cues remain ill-defined. The rapid development of single molecule and single cell analyses should allow us to now define exactly how chromatin remodellers, signalling molecules and transcription factors all interact at regulatory sequences to allow cells to respond quickly to changes in the local environment.

## Materials and Methods

### Cell lines and culture conditions

hiPSCs were a generous gift of Prof. Austin Smith (Takashima et al., 2014). Endogenously tagged MBD3-3xFLAG hiPSCs were made using a CRISPR/Cas9 gene editing approach to insert 3xFLAG immediately upstream of the MBD3 stop codon using a guide RNA targeting the sequence 5’-GAGCGAGTGTAGCACAGGTG-3’ (Supplemental Fig. 1). *MBD3*-KO cells were generated replacing exons 2 and 3 with a puromycin resistance cassette using CRISPR/Cas9-mediated targeting and guide RNAs targeting the sequences 5’-GGCGGTGGACCAGCCGCGCC-3’ and 5’-GTCGCTCTTGACCTTGTTGC-3’. A correctly targeted heterozygous clone was then transiently transfected with Dre recombinase prior to a second round of targeting to generate a homozygous null line (Supplemental Fig. 1). The MBD3 Rescue line was made by transfecting the *MBD3*-KO iPS line with a construct containing a CAG promoter driving expression of full-length MBD3-3xFLAG, followed by an IRES and a hygromycin resistance gene, and a polyA sequence from the human *PGK* gene. Hygromycin resistant cells were expanded and tested for MBD3-3xFLAG expression (e.g. Figure 2A).

hiPSCs were cultured in mTESR1 (StemCell Technologies) media or E8 medium (made in house, prepared according to (Chen et al., 2011)) on vitronectin coated plates. hiPSCs were passaged using an enzyme-free passaging reagent (ReleSR, StemCell Technologies) and plated as small clumps.

Neuroectoderm differentiation was induced based on (Vallier, Touboul et al., 2009). hiPSCs were plated as clumps (day −1) in chemically defined medium with Polyvinyl Alcohol (CDM-PVA) supplemented with hActivin A (10ng/µl) and FGF2 (12ng/µl) on 0.1% gelatin coated plates pre-treated overnight at 37°C with MEF media (Advanced DMEM-F12, 10% FBS, 2 mM L-glutamine, 1x penicillin/streptomycin). The original composition of CDM is 50% IMDM (Gibco) plus 50% F12 Nutrient-MIX (Gibco), supplemented with 4 ug/ml of insulin (Roche), 15 µg/ml transferrin (Roche), 450 µM monothioglycerol (Sigma), Chemically Defined lipid concentrate (Invitrogen). The next day (day 0), hiPSCs were cultured in CDM-PVA supplemented with SB431542 (10 µM, Tocris), FGF2 (12 ng/ml, R and D Systems) and Noggin (15 ng/ml, Peprotech) for 12 additional days. The cells were harvested using Accutase at 3, 6 and 12 days. The media was changed every day.

Definitive endoderm differentiation was induced according to (Yiangou, Grandy et al., 2019). Cells were cultured in CDM-PVA supplemented with 100 ng/ml Activin A (produced in house), 80 ng/ml FGF2 (produced in house), 10 ng/ml BMP4 (R&D Systems), 10 µM LY294002 (Promega) and 3uM CHIR99021 for one day, with CHIR99021 omitted on the second day. From day three onwards, cells were cultured in RPMI basal medium, supplemented with 100 ng/ml Activin A and 80 ng/ml FGF2 on day 3. From day 4 onwards, RPMI was supplemented with 50 ng/ml Activin A only. The cells were harvested using Accutase at days 2, 4, 6 and 8. The media was changed every day.

mEpiSCs were derived from *Mbd3*^*Flox/Δ*^ mESCs and subsequently transiently transfected with Cre recombinase to create *Mbd3*^*Δ/Δ*^ cells. *Mbd3*^*Δ/ Δ*^ mEpiSCs were independently derived from *Mbd3*^*Δ/Δ*^ ES cells. mEpiSC cultures were maintained in N2B27 supplemented with FGF2 (12 ng/µl), Activin A (20 ng/µl), XAV939 (2 mM, Sigma) on fibronectin (15 µg/ml) pre-coated plates. The cells were harvested using Accutase at 2, 4 and 8 days. The media was changed every day. For neural differentiation cells were plated on laminin-coated plates in N2B27 containing 1 µM A83-01 (StemMACS).

### Gene expression analysis

was carried out as described (Burgold et al., 2019). Briefly, total RNA was isolated using RNA mini easy kit (Qiagen) and reverse transcribed using random hexamers and Superscript IV Reverse Transcriptase (Invitrogen). Quantitative PCR was carried out using gene-specific primers and Sybrgreen incorporation, or Taqman reagents on a StepOne or ViiA7 real time PCR system (both Applied Biosystems).

### TAQMAN PROBES

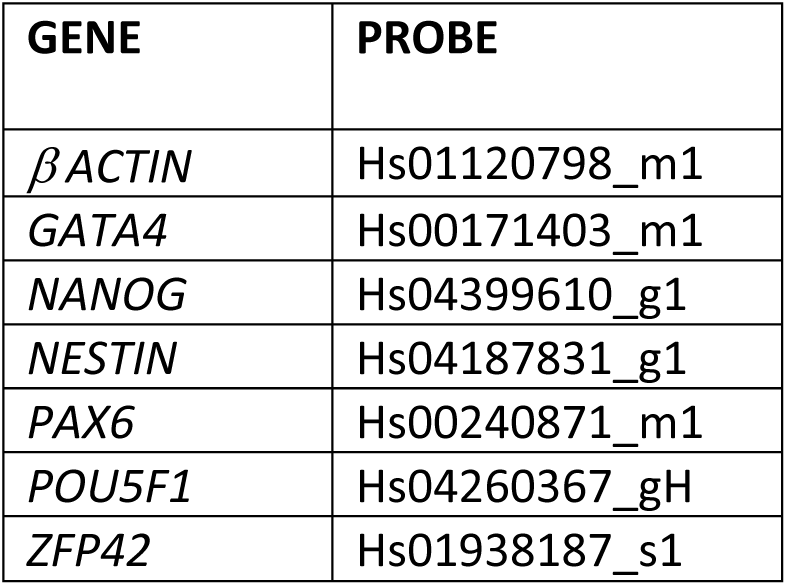

### PRIMERS

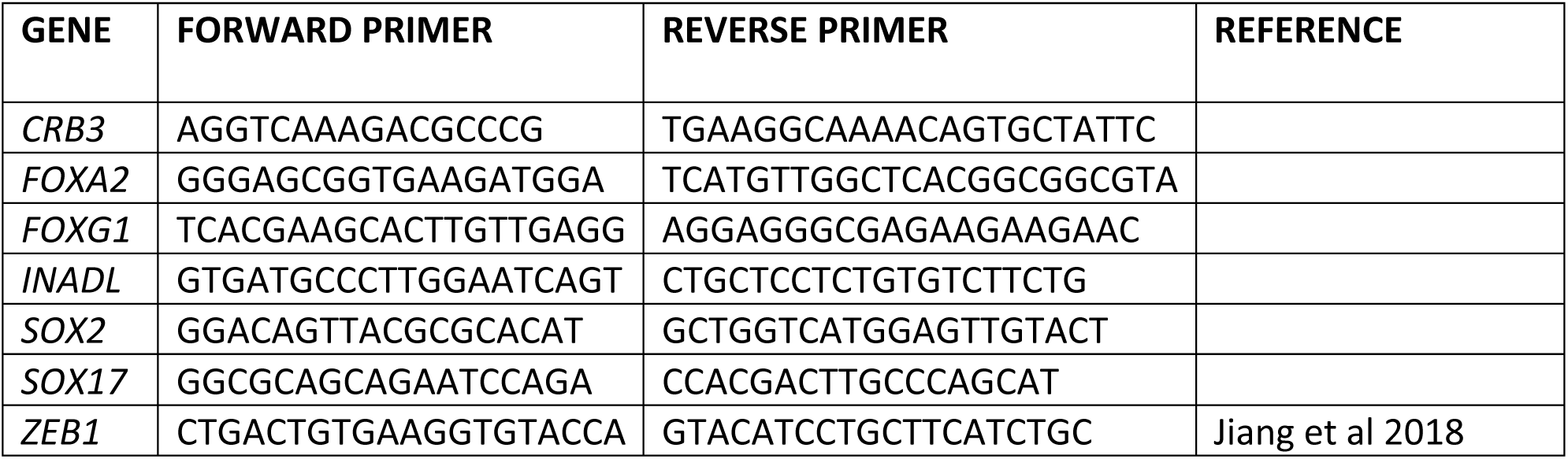

### Nuclear extraction, immunoprecipitation and Proteomics

Nuclear extraction was an immunoprecipitation was performed as described (Burgold et al., 2019). Original western blot images are available on Mendeley Data: XXXX. Antibodies used in this study are indicated below:

| ANTIBODY | RAISED IN | COMPANY | CATALOGUE NUMBER | DILUTION FOR WESTERN BLOT |
| --- | --- | --- | --- | --- |
| αCHD4 | Mouse | Abcam | ab70469 | 1:5000 |
| αGATAD2B | Rabbit | Bethyl Labs | A301-281A | 1:2000 |
| αLAMIN B1 | Rabbit | Abcam | ab133741 | 1:10 000 |
| αMBD3 | Rabbit | Abcam | ab157464 | 1:5000 |
| αMTA2 | Mouse | Abcam | ab50209 | 1:5000 |

Mass spectrometry was carried out as described (Burgold et al., 2019, Kloet et al., 2018, Smits, Jansen et al., 2013). Briefly, nuclear extract was prepared from a human iPS cell line in which a 3xFLAG tag was knocked in to the endogenous *MBD3* locus, or from two independent mouse epiStem cell lines similarly modified as described (Burgold et al., 2019). One preparation of nuclear extract from each cell line was divided into thirds, which were independently processed for proteomic analyses. Proteins associated with 3xFLAG-tagged MBD3 were purified using anti-FLAG sepharose (Sigma) and processed for mass spectrometry as described (Smits et al., 2013). The resulting data were processed as in (Kloet et al., 2018).

### RNA-seq and analysis

Sequencing libraries were prepared using the NEXTflex Rapid Directional RNA-seq kit (Illumina) or SMARTer® Stranded Total RNA-Seq Kit v2—Pico Input Mammalian (Takara Bio) and sequenced on the Illumina platform at the CRUK Cambridge Institute Genomics Core facility (Cambridge, UK). Illumina sequence files were converted into FASTQ format. The short sequence reads (75 nucleotides) were aligned to the Human reference genome (hg38; http://genome.ucsc.edu/) or to the Mouse reference genome (mm10; http://genome.ucsc.edu/) and assigned to genes using BWA (Li & Durbin, 2009). We used the Subread package (R statistical tool; http://www.r-project.org/) to count aligned reads. Differentially expressed genes were identified using R package edgeR (Chen, Lun, & Smyth, 2016). We used no fold change filtering and results were corrected for multi-testing by the method of the False Discovery Rate (FDR) at the 1% level. Differentially expressed genes were clustered using the unsupervised classification method of the Kmeans (Soukas, Cohen, Socci, & Friedman, 2000). Heat maps were done using the pheatmap function (R statistical tool; http://www.r-project.org/). Functional annotation enrichment for Gene Ontology (GO) terms were realised using HumanMine [http://www.humanmine.org] (Smith, et al., 2012) or MouseMine database [http://www.mousemine.org]. Benjamini-Hochberg corrected P values of less than 0.01 were considered significant. GO terms were submitted to REVIGO, a web server that takes long lists of GO terms and summarizes them in categories and clusters of differentially expressed genes by removing redundant entries (Supek, Bošnjak, Škunca, & Šmuc, 2011). We used i-*cis*Target tool (Imrichová, Hulselmans, Kalender Atak, Potier, & Aerts, 2015) to look for enrichment in TF position weight matrices and potential binding sites in the regulatory regions of co-expressed genes. i-cisTargetX computes statistical over representation of DNA motifs and ChIP-seq peaks in the non-coding DNA around sets of genes. The enrichment was considered significant when the Normalized Enrichment Score (NES) was higher than 5.

### Data Availability

RNA-seq data are available with the Array Express accession number E-MTAB-8753. The mass spectrometry proteomics data have been deposited to the ProteomeXchange Consortium via the PRIDE partner repository with the dataset identifier PXD016967. All original western blot images are available at Mendeley Data: DOI: 10.17632/4t99j4c7gx.1.

## Acknowledgments

We thank Austin Smith for providing the hiPSC line, Sabine Dietmann, Maike Paramor, Vicki Murray, Peter Humphreys and Sally Lees for technical assistance and advice, Sabine Dietmann, Denis Seyres and Laurence Röder for comments on the manuscript and members of 4DCellFate Consortium for useful discussions and suggestions. Funding to the BH and MV labs was provided through EU FP7 Integrated Project “4DCellFate” (277899). The BH lab further benefitted from grants from the Medical Research Council (MR/R009759/1), the Wellcome Trust (206291/Z/17/Z) and the Isaac Newton Trust (17.24(aa)), and core funding to the Cambridge Stem Cell Institute from the Wellcome Trust and Medical Research Council (203151/Z/16/Z). The Vermeulen lab is part of the Oncode Institute, which is partly funded by the Dutch Cancer Society (KWF).

## Author Contributions

NR and BH devised the study; SG, JC, OO, SK, TB, NR and BH generated the data; RR analysed high throughput sequencing data, SK and MV generated and analysed proteomics data and RR and BH wrote the manuscript with input from other authors.

---

**Figure S1.**
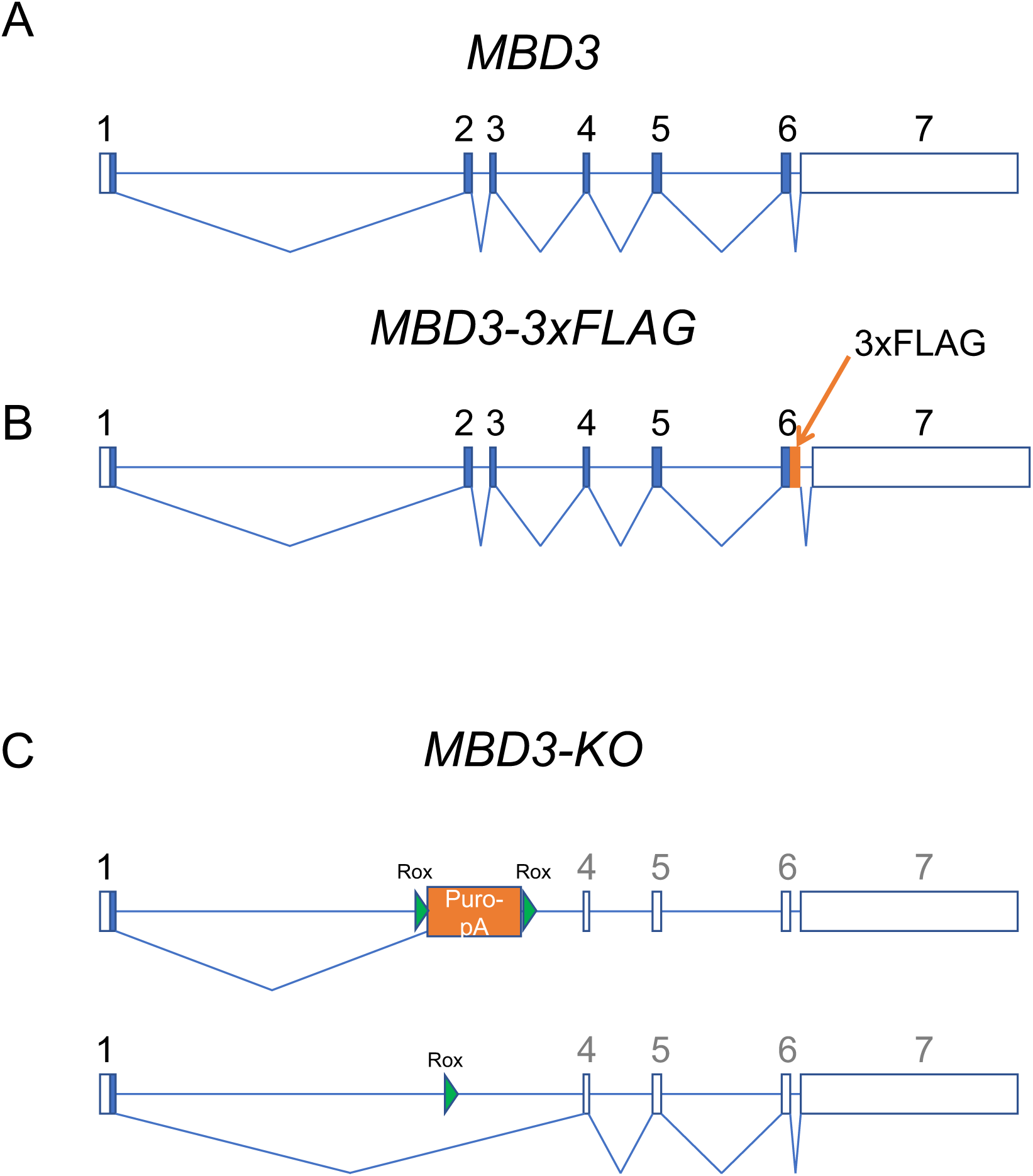
Human alleles used in this study. A) Schematic of the human *MBD3* locus. Filled boxes represent coding exons, open boxes represent non-coding exons. B). Schematic of the *MBD3-FLAG* allele. A 3xFLAG epitope was knocked-in immediately upstream of the stop codon in Exon 6 on one *MBD3* allele. C). Schematic of the *MBD3*-KO alleles. Exons 2 and 3 were replaced by a Puromycin resistance cassette flanked by Rox sites. After the first round of targeting the Puromycin cassette was removed by transient Dre expression, followed by a second round of targeting to mutate both *MBD3* alleles.

**Figure S2.**
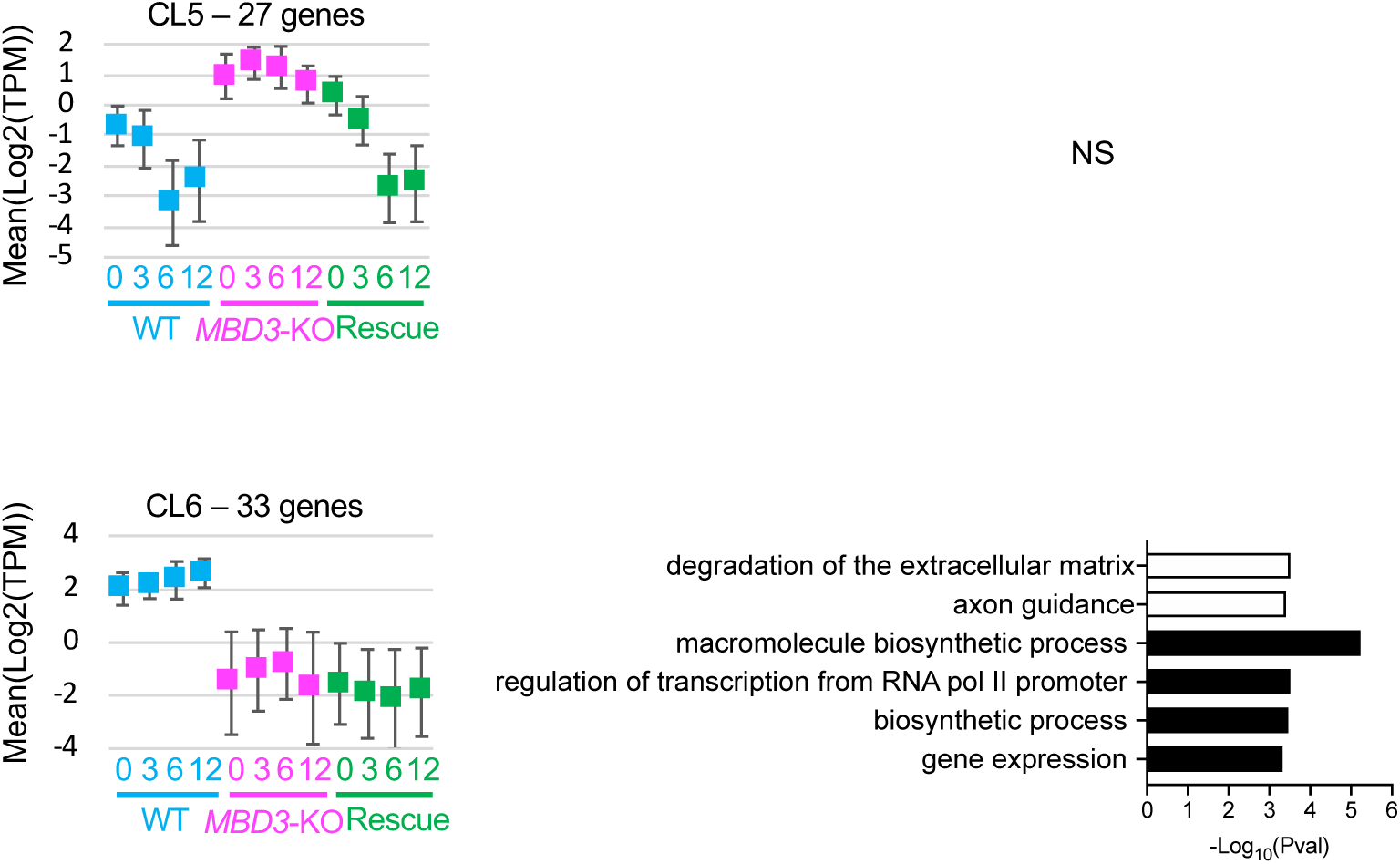
Expression and GO-term plots for Clusters 5 and 6 from Figure 4A. There were no significant (P.adj≤0.01) GO terms associated with Cluster 5 genes. Error bars indicate standard deviations of average expression.

**Figure S3.**
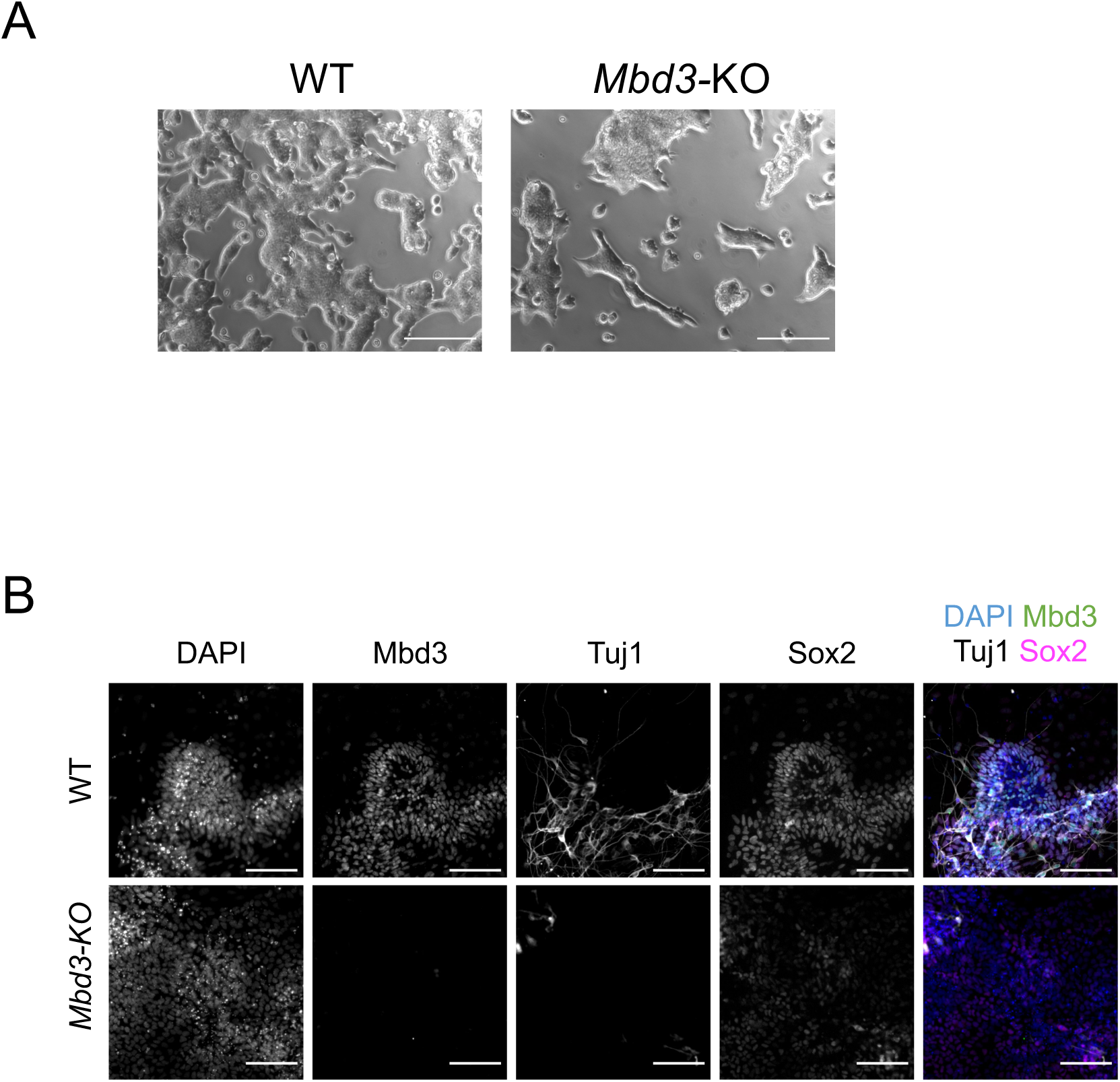
Mouse *Mbd3*-KO epiStem cells. A) Phase contrast images of wild type or *Mbd3*-KO mEpiSCs in self-renewing conditions. B). Wild type or *Mbd3*-KO cells after 8 days of neural differentiation were stained with antibodies indicated at top or DAPI. The final panels show merged images. Scale bars represent 100µm.

**Figure S4.**
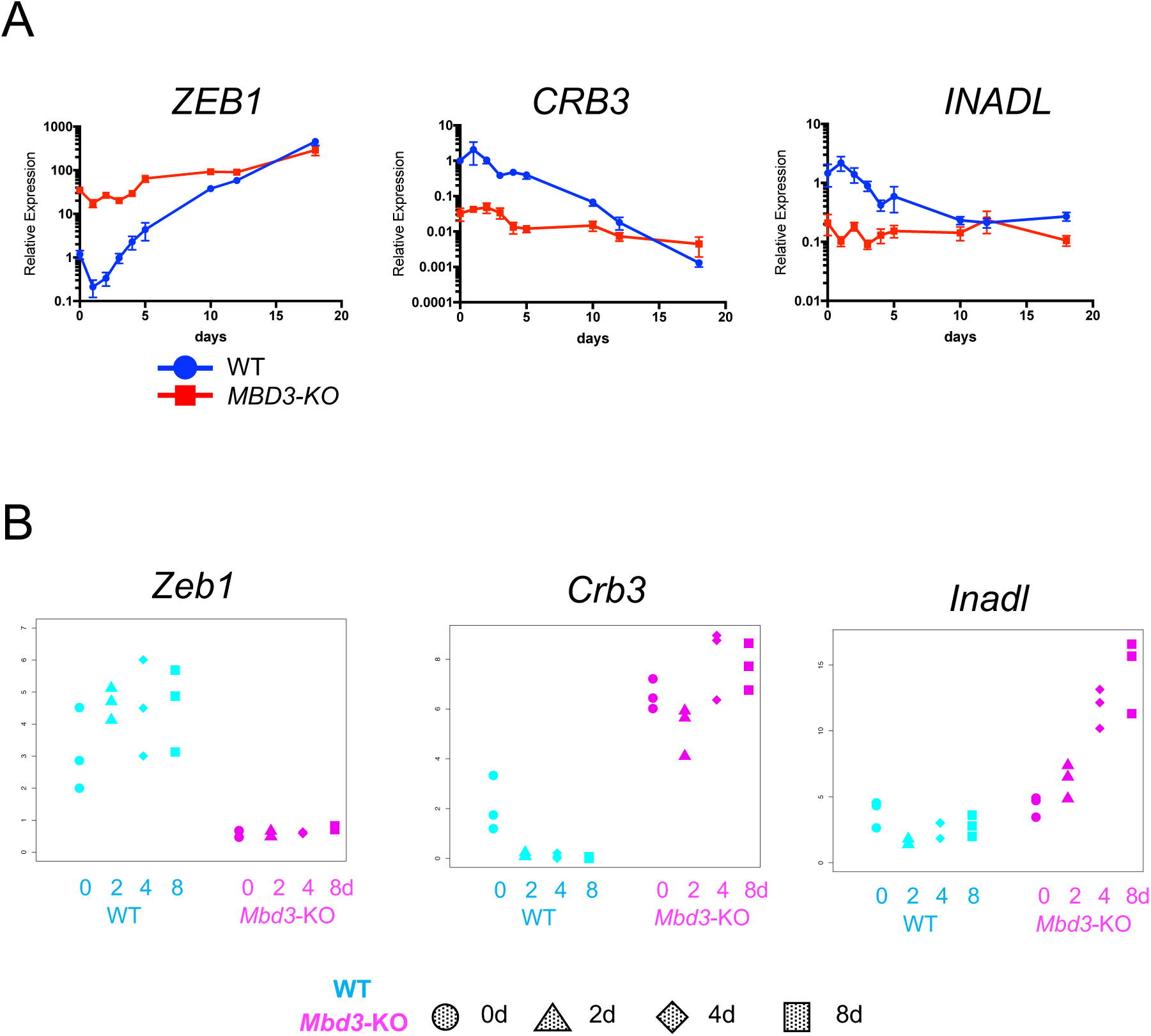
Expression profiles of *ZEB1, CRB3* and *INADL* in hiPSCs and mEpiSCs. A) RT-qPCR of *ZEB1, CRB3* and *INADL* in hiPSCs during neuroectodermal differentiation. The y-axis shows the relative expression and the x-axis shows the day of differentiation. The blue line represents the WT while the red line represents *MBD3*-KO hiPSCs (n=3 replicates/timepoint/condition). Error bars indicate the standard deviation. B) Expression profiles of *Zeb1, Crb3* and *Inadl* from mEpiSC RNA-seq data. The y-axis represents TPM (Transcript Per Million) while the x-axis and the different shapes represent the days of differentiation (n≥3 replicates/timepoint/condition). Blue indicate the WT and magenta represent Mbd3-KO mEpiSC.

